# SCIM: Universal Single-Cell Matching with Unpaired Feature Sets

**DOI:** 10.1101/2020.06.11.146845

**Authors:** Stefan G. Stark, Joanna Ficek, Francesco Locatello, Ximena Bonilla, Stéphane Chevrier, Franziska Singer, Tumor Profiler Consortium, Gunnar Rätsch, Kjong-Van Lehmann

## Abstract

**Motivation:** Recent technological advances have led to an increase in the production and availability of single-cell data. The ability to integrate a set of multi-technology measurements would allow the identification of biologically or clinically meaningful observations through the unification of the perspectives afforded by each technology. In most cases, however, profiling technologies consume the used cells and thus pairwise correspondences between datasets are lost. Due to the sheer size single-cell datasets can acquire, scalable algorithms that are able to universally match single-cell measurements carried out in one cell to its corresponding sibling in another technology are needed.

**Results:** We propose Single-Cell data Integration via Matching (SCIM), a scalable approach to recover such correspondences in two or more technologies. SCIM assumes that cells share a common (low-dimensional) underlying structure and that the underlying cell distribution is approximately constant across technologies. It constructs a technology-invariant latent space using an auto-encoder framework with an adversarial objective. Multi-modal datasets are integrated by pairing cells across technologies using a bipartite matching scheme that operates on the low-dimensional latent representations. We evaluate SCIM on a simulated cellular branching process and show that the cell-to-cell matches derived by SCIM reflect the same pseudotime on the simulated dataset. Moreover, we apply our method to two real-world scenarios, a melanoma tumor sample and a human bone marrow sample, where we pair cells from a scRNA dataset to their sibling cells in a CyTOF dataset achieving 93% and 84% cell-matching accuracy for each one of the samples respectively.

**Availability:** https://github.com/ratschlab/scim

## 1 Introduction

The ability to dissect a tissue into its cellular components to study them individually or to investigate the interplay between the different cell-type fractions is an exciting new possibility in biological research that has already yielded important insights into the dynamics of various diseases including cancer (Tirosh *et al.*, 2016; Chevrier *et al.*, 2017). Recent advances in single-cell technologies enable molecular profiling of samples with greater granularity at the transcriptomic, proteomic, genomic as well as the functional assays level (Rozenblatt-Rosen *et al.*, 2017; Irmisch *et al.*, 2020). Each data modality produces different types and levels of information that need to be integrated and related to one another to truly grasp the mechanisms at play in the tissue microenvironment and to obtain a more comprehensive molecular understanding of the studied sample. Although technologies capable of measuring two modalities simultaneously are emerging (Stoeckius *et al.*, 2017; Zhu *et al.*, 2020), their scalability and widespread use are still limited. While multiple data integration tools have been developed recently, most approaches either depend on feature correspondences (Stuart *et al.*, 2019; Welch *et al.*, 2019) or are limited to a specific input type, for instance, scRNA and scDNA data (Campbell *et al.*, 2019; McCarthy *et al.*, 2020). To the best of our knowledge only two other approaches have been published (Amodio and Krishnaswamy, 2018; Welch *et al.*, 2017) with similar capabilities to SCIM. MAGAN (Amodio and Krishnaswamy, 2018) is a Generative Adversarial Network (GAN) capable of aligning the manifold between two technologies that relies on a feature correspondence loss. MATCHER (Welch *et al.*, 2017) is based on a gaussian process latent variable model (GPLVM) (Lawrence, 2004) that can integrate technologies if their underlying latent structures can be represented in one dimension, applicable, for example, to model monotonic temporal processes. Other yet unpublished methods, such as MMD-MA (Liu *et al.*, 2019) and UnionCom (Cao *et al.*, 2020), rely on large kernel matrices which limit their scalability when using datasets of the sizes generally produced by molecular profiling.

Here, we propose SCIM, a method to match cells across different single-cell ‘omics technologies. Our approach consists of two parts. First, we build an integrated latent space where representations are invariant to their corresponding technologies inspired by a model proposed previously (Yang and Uhler, 2019) and further extended in Yang *et al.* (2019). Then, we apply a cell-to-cell matching strategy that efficiently extracts cross-technology cell matches from the latent space. SCIM assumes a shared latent representation between technologies but, unlike other approaches, does not require one-to-one or overlapping correspondences between feature sets. Individual technologies often consume samples thus, the input material provided to each profiling approach typically is an aliquot from a common sample cell suspension. Notwithstanding, given that the technology-specific datasets come from the same sample, (i.e., cell mix), expecting the same underlying distribution is an appropriate assumption. SCIM scales well in the number of cells in the input through the use of neural-nets, end-to-end training, and an efficient bipartite matching algorithm. The training scheme allows for the addition of an arbitrary number of technologies, which can be trained in parallel (see **Figure 1**).

**Figure 1:**
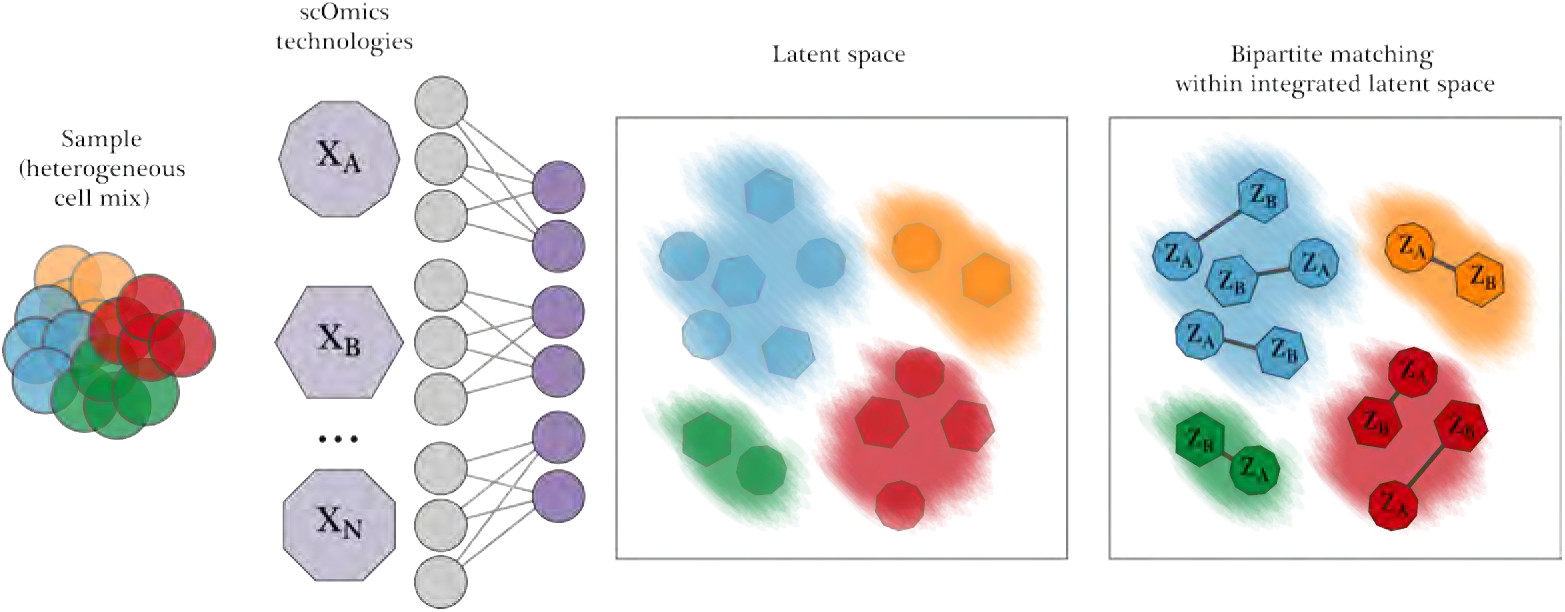
SCIM performs a pairwise matching of cell across multiple single-cell ‘omics technologies. We assume that the input of each technology comes from the same (or similar) heterogeneous cell mix, depicted on the left. Technologies generate a set of single cell ‘omics datasets (violet polygons) in parallel (e.g. *X*_*A*_, *X*_*B*_, *X*_*N*_). SCIM proceeds to map cells into a technology-invariant latent space (left box) using an autoencoder framework and an adversarial term to keep technologies well integrated. Here the latent representations capture the underlying structure in the cell mix (colored clouds) and analogous codes from different technologies (colored polygons) are placed in proximity. To integrate datasets, a fast bipartite matching scheme is applied, matching cells pairwise among datasets to cross-technology analogs, using their latent representations (right box).

## 2 Methods

SCIM matches cells from a source technology to cells in one or multiple target technologies in two main steps. First, an integrated, technology-invariant latent space is produced using an encoder/decoder framework based on Yang and Uhler (2019). Then, cells are paired across different technologies via their latent representations using a version of the fast bipartite matching algorithm.

### 2.1 Model

Autoencoders produce low dimensional representations of data by learning a pair of encoder and decoder functions, with parameters *ϕ* and *ψ* respectively. The encoder maps input data into a lower-dimensional space, called the latent space, while the decoder tries to reconstruct the input data from its latent representation. The popular Variational Autoencoders (VAEs) take a generative approach to this problem (Kingma and Welling, 2013). Here *ϕ* parameterizes the likelihood of the data given the latent representation *p*_*ϕ*_(*x* | *z*) and *ψ* parameterizes the posterior probability of its latent representation *q*_*ψ*_(*z* | *x*). VAEs jointly learn *ϕ* and *ψ* to maximize a lower bound to the probability of the data *p*(*x*; *ϕ, ψ*), achieved in practice by minimizing

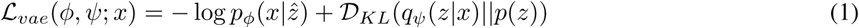

where 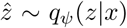, 𝒟_*KL*_ is the Kullback-Leibler (KL) divergence, and *p*(*z*) is a prior distribution over latent representations. Often *p*(*z*) and *q*_*ψ*_(*z* | *x*) are restricted to Gaussian forms since the KL divergence then has a closed-form solution.

#### 2.1.1 Constructing a Technology-Invariant Latent Space

SCIM encodes datasets into a shared latent space, which has ideally two properties. As in the VAE, inputs should be able to be reconstructed from their latent representations. Additionally, the latent representations of each technology should be integrated well such that they are indistinguishable from each other. In a successful integration the resulting latent space will have corresponding cells across all technologies represented in close proximity.

To construct an integrated latent space, SCIM employs the following networks: a pair of encoder (*ϕ*_*k*_) and decoder (*ψ*_*k*_) networks for each technology *k* and a single discriminator network (*γ*) acting on the latent space. The discriminator is a binary classifier trained to identify the latent representation of a source technology from latent representations of all other technologies using a binary cross entropy loss.

SCIM yields an integrated latent space by minimizing the reconstruction error while adversarially fooling the discriminator. For notational brevity we now let *ϕ*_*k*_ and *ψ*_*k*_ also represent the probability distributions they parameterize. Given the measurements of a batch of cells from the target technology, *x*_*t*_, and the (fixed) latent representations of a batch of cells from the source technology, 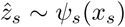, SCIM minimizes the following objective

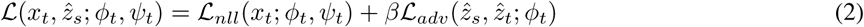

ℒ _*nll*_(*x*_*t*_; *ϕ*_*t*_, *ψ*_*t*_) is the negative log-likelihood of the inputs under their reconstruction. ℒ _*adv*_ is the discriminator’s classification error when trying to classify the latent representation samples 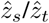 as the target/source technology. *β* is a hyperparameter weighing the influence of the adversarial loss. At the same time, *γ* is trained to correctly classify the technology of the 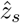 and 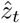 samples.

More intuitively, this framework can be seen as learning a VAE on each technology where the prior distribution is defined by the latent representations of the other technologies. ℒ_*adv*_ can be interpreted as a divergence measure where, through the use of adversarial techniques, samples may be used in lieu of their potentially intractable probability distributions. Thus, the framework is equivalent to a set of Adversarial Autoencoders (Makhzani *et al.*, 2015) or Wasserstein Autoencoders (Tolstikhin *et al.*, 2017) which share a single discriminator.

#### 2.1.2 The Orientation of Latent Space

Correctly orienting the latent space in an unsupervised manner is a challenging task (Locatello *et al.*, 2018; Yang and Uhler, 2019). Consider, for example, a simple monotonic temporal process. The latent representations for one dataset could be oriented from start to end, while another could be oriented from finish to start (Welch *et al.*, 2017). Equation 2 is satisfied, the representations are well integrated and inputs can be correctly reconstructed from them, yet the inter-dataset relationships are misaligned.

Makhzani *et al.* (2015) address a similar problem by concatenating one-hot representations of labels reflecting intra-technology structure (e.g. cell type is an appropriate choice for ‘omics datasets) to the discriminator inputs, showing that this supervision is necessary to orient the latent space. Recently, Locatello *et al.* (2019) argued that only a small number of labels are actually needed to achieve orientation. To this end, we adopt a semi-supervised approach by adding a “censored” label and randomly relabel cells in the training set.

#### 2.1.3 Model Architecture

Unless specified otherwise, we adopt the following architecture settings. All networks use the ReLU activation. We set the latent dimension of all models to 8, but observed this choice to be flexible. We use discriminator networks with 2 layers and 8 hidden units each. The Spectral Normalization framework (Miyato *et al.*, 2018) is used during training, which has been argued to stabilize discriminator training by effectively bounding its gradients. We use a Gaussian activation for all decoders, a 2 layer architecture with 64 hidden units for all simulated data networks, a 2 layer architecture with 8 hidden units for all CyTOF networks, and a 2 layer architecture with 64 hidden units for all scRNA networks. The number of features and complexity of data is considered when choosing capacity and depth.

#### 2.1.4 Optimization

Optimization proceeds by iteratively fixing one technology as the source and one technology as the target. In the case of more than two technologies, the technology corresponding to the discriminator’s positive class must either be the source or target technology. The codes of the source technology are fixed and Equation 2 is minimized with gradient updates to the encoder and decoder, *ϕ*_*t*_ and *ψ*_*t*_, of the target technology using gradients computed on the batch *x*_*t*_. After each update, the discriminator is trained to correctly classify 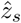 and 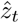. All networks in SCIM are optimized using the ADAM algorithm (Kingma and Ba, 2014).

We initialize SCIM by first training a VAE (Kingma and Welling, 2013) on a single source technology, and use the latent representations as the first set of 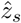. Unless specified otherwise the VAE is trained for 256 epochs using *β* = 0.01 and a learning rate of 0.0005. A small value of *β* is needed for structure to be retained in the latent representations.

#### 2.1.5 Latent Space Evaluation and Model Selection

Due to the min-max nature of adversarial training, model comparison is challenging since one cannot directly compare the minimized objective functions of converged models (Lucic *et al.*, 2017). The computer vision community has introduced a number of metrics specific to the image domain to help compare models (Heusel *et al.*, 2017; Salimans *et al.*, 2016). Here we need to validate the quality of a set of lower dimensional latent representations.

Therefore we use a k-Nearest Neighbor (kNN) based divergence estimator (Wang *et al.*, 2009) to quantitatively evaluate the quality of the integrated latent space. The divergence score between two sets of codes *Z*_*s*_ and *Z*_*t*_ is calculated as:

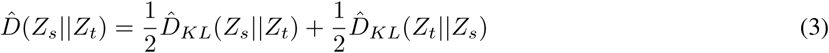

Where

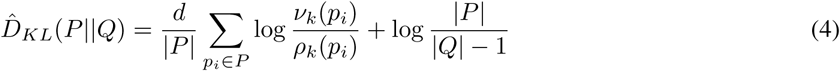

where *ν*_*k*_(*p*_*i*_) and *ρ*_*k*_(*p*_*i*_) are the distances from *p*_*i*_ to the *k*^*th*^ nearest neighbor in the sets *P* and *Q* respectively and all *p*_*i*_ ∈ ℛ^*d*^. This estimator approximates a symmetric variant of a Kullback-Leibler (KL) divergence, a measure of how much two distributions differ, using only empirical data. The divergence estimate is computed between the latent representations of the source technology and the target technology.

This approach draws inspiration from a proposed framework from (Yang and Uhler, 2019), where expression profiles are decoded from the latent space. We were able to utilize a kNN based divergence estimator (Wang *et al.*, 2019), to address typical problems in adversarial training. Further, SCIM does not decode values from latent space, but the low-dimensional representation is used solely to match the cells. Thus, the true observed marker abundances per cell pair, measured with different technologies, can be used for any downstream analysis. Moreover the latent space matching matching may compensate for sub-optimal integration, providing an additional advantage over bare decoding.

### 2.2 Bipartite Matching of Latent Representations

The obtained shared latent representation can be used for finding corresponding cells across technologies. Each cell is now characterized by a low-dimensional vector of latent codes, which are in one-to-one correspondence across technologies. We first represent the data as a graph, where the nodes correspond to cells and edge weights correspond to the Euclidean distances in latent space between the cells. To find the best pairwise matching efficiently, we phrase the task as a combinatorial bipartite matching problem (Ahuja *et al.*, 1993; Dell’Amico and Toth, 2000). In other words we identify edges connecting the cells that would result in a minimal total cost of all matches. In order to achieve this, we build a k-Nearest Neighbors (kNN) graph to identify a set of potential matches and reduce the complexity of the problem. Then, we extend the graph to account for single-cell data characteristics and solve the bipartite matching within a general framework of Minimum-Cost Maximum-Flow problems (Ahuja *et al.*, 1993; Klein, 1967).

#### 2.2.1 k-Nearest Neighbor Approximation

Given the large number of cells profiled in single-cell data, we reduce the search space to the *k* most likely potential matches. Two kNN graphs are built: 1) using source data queried by the target technology cells, and 2) using target data queried by the source technology cells. A union of the established connections is used for further analysis. The sparsity of connections, regulated by the choice of hyperparameter *k*, corresponds to the trade-off between the computational performance (memory usage, run time) and the matching accuracy.

#### 2.2.2 Bipartite Matching via Minimum-Cost Maximum-Flow

Based on a cost matrix, we aim at finding the maximum number of cell pairs with minimum cost. This corresponds to finding a maximum flow that can be pushed through the graph, where each edge between cells has capacity 1, while minimizing the overall cost. To solve the Minimum-Cost Maximum-Flow problem in a computationally efficient way we use an implementation of the network simplex algorithm (Király and Kovács, 2012).

#### 2.2.3 Relaxation of One-to-One Matching by Graph Extensions

Bipartite matching approach makes the assumption that each cell has one and only one direct corresponding sibling in the other technology. To allow for mismatches due to expected variation in cellular composition, we expand the kNN graph with sparse connections by adding a densely connected *null* node with high capacity and high assignment cost. This allows to capture potentially poorly matched cells. The magnitude of the null match penalty corresponds to a given percentile *p* of the overall costs and is a hyperparameter. The extended graph structure is depicted in **Figure 2**, where *R* and *S* refer to the root and sink nodes, respectively. Furthermore, to account for differences in the number of cells between modalities (*n, m*), we allow for one-to-many matches by increasing the capacity of the edges incoming to the sink (*u*_*i*_ for *i* ∈ {1, *…, m*}), assuming the nodes linked to the sink correspond to the smaller dataset (*m* ≤ *n*). To prevent all matches from collapsing onto a very small set of nodes, we constrain the sum of incoming sink capacities, excluding the null node, to equal the cardinality of the bigger dataset (*n*).

**Figure 2:**
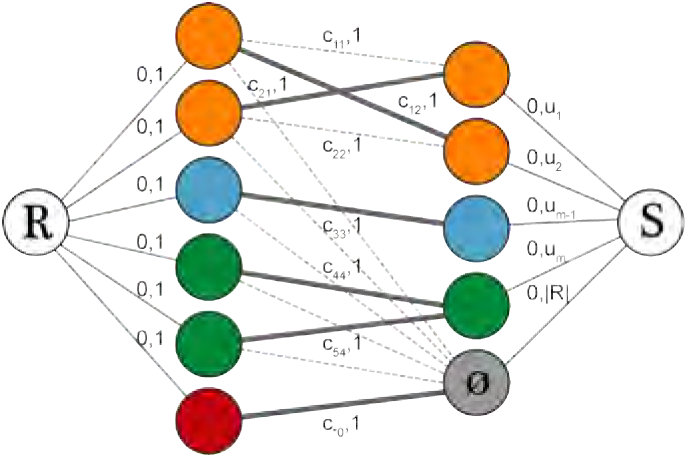
Fast bipartite matching using a customized Minimum-Cost Maximum-Flow framework. Nodes correspond to cells and edges to the sparse connections between them, resulting from a kNN search. *R* and *S* represent root and sink nodes. Edge labels (a, b) indicate matching cost and edge capacity, respectively. Many-to-one matches in unbalanced datasets are enabled by increasing the capacities *u*_*i*_ (for *i* ∈ 1, *…, m*). The *null* node, colored in grey, captures matches of cells (from the bigger dataset on the left-hand side of the graph) that lack a close enough analog in the other technology. Its capacity equals the cardinality of the bigger dataset and the cost *c*_*0_, i.e., null match penalty, is relatively high. The thicker lines linking the nodes represent the actual matches selected by the algorithm.

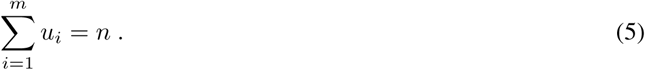

The capacities are distributed uniformly across the sink edges. In case of a high disproportion in the number of cells profiled with different technologies, the penalty and uniformity of capacities may limit the total number of matches. Therefore, in such cases the capacity of the sink edges remains unbounded and is set to infinity. In case more than two data modalities are present, the bipartite matching is solved sequentially by obtaining pairwise matches between technologies.

#### 2.2.4 Matching Evaluation

The quality of matching is evaluated on different levels. First, the accuracy corresponding to the fraction of true positives with regards to cell-type label is reported. Cell-types can be determined in a technology specific manner and the accuracy is reported on a common denominator. If more fine-grained cellular information is available, such as pseudotime, a direct comparison of this quantity is carried out. Furthermore, in real-world data settings we utilize the raw marker expression to investigate correspondence of the matched cells. Namely, Spearman’s and Pearson’s correlation coefficients are computed between the expression values across matches.

## 3 Data

### 3.1 Simulated Data

Using PROSSTT (Papadopoulos *et al.*, 2019), we generate three single-cell ‘omics-styled technologies which share a common latent structure without direct feature correspondences. PROSSTT parameterizes a negative binomial distribution given a tree representing an underlying temporal branching process. By using the same tree and running PROSSTT under different seeds, we obtain three datasets with a common latent structure yet lacking any correspondences between features. We used a five branch tree with different branch lengths (**Figure 3**). Each dataset contains 64,000 cells with 256 markers.

**Figure 3:**
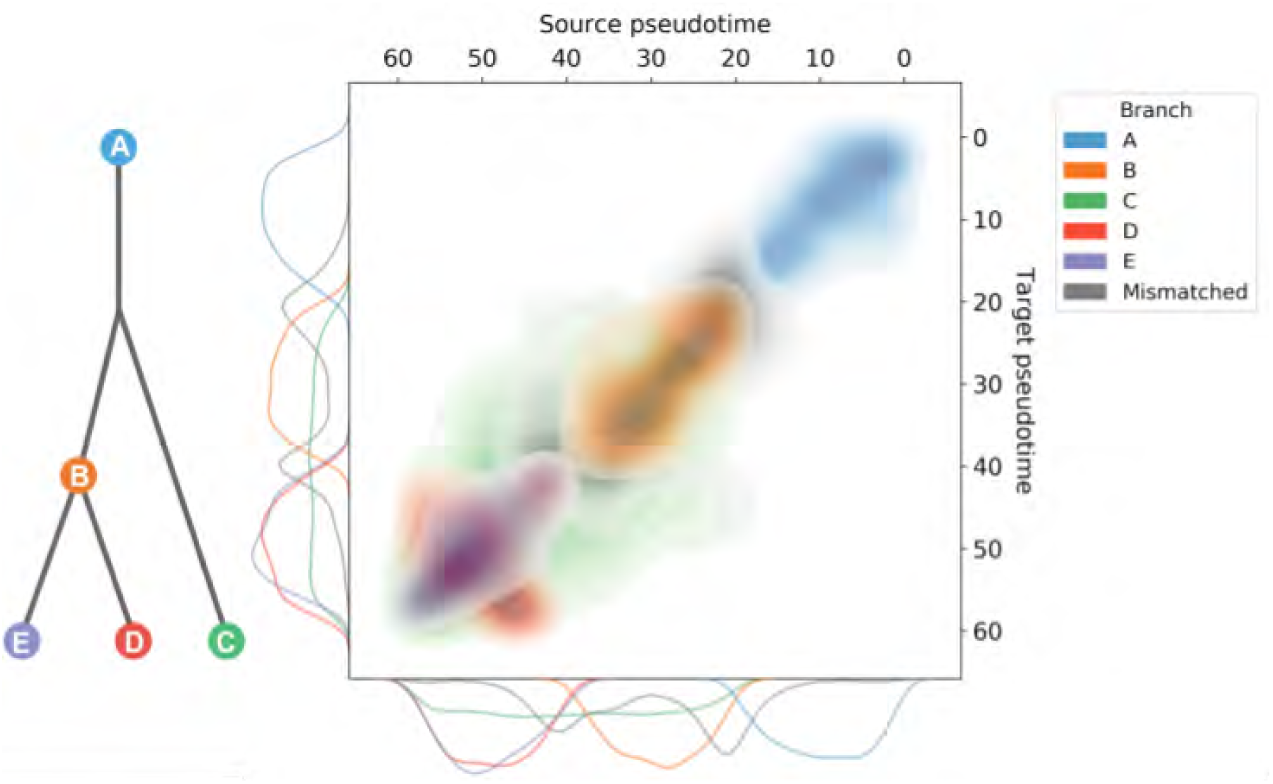
Evaluation of cross-technology cell matches made by SCIM on the simulated data. Cells are matched across datasets pairwise using the bipartite matching scheme. The tree defining the temporal branching process is shown on the left and the density plot for matched pseudotime values between the source technology and the target technology, on the right. Cells matched to the same branch label are colored (accuracy: 86%), while mismatches are depicted in grey. Marginal distributions of cell pseudotime for each branch are shown at the bottom and left of the density plot. We report a correlation of 0.83 (Spearman) and 0.86 (Pearson) for pseudotime label pairs.

### 3.2 Single-cell Profile of a Melanoma Patient

The motivating dataset for our research questions is generated by the Tumor Profiler (TuPro) Consortium (Irmisch *et al.*, 2020) as part of a multi-center, multi-cancer study comprising metastatic tumors from a cohort of deeply phenotyped individuals. Each patient’s data is analyzed with multiple technologies, including scRNA-sequencing (Tang *et al.*, 2009), Cytometry by Time Of Flight (Bandura *et al.*, 2009, CyTOF), all capable of dissecting the tumor microenvironment and providing single-cell level, complementary information about the sample of interest. Although cell identity is lost throughout the process, the cells investigated by both technologies stem from the same population (i.e., were obtained from an aliquot of a common cell suspension).

#### 3.2.1 CyTOF Data Preparation

The patient’s sample was profiled with CyTOF using a 40-markers panel designed for an in-depth characterization of the immune compartment of a sample. Data preprocessing was performed following the workflow described in (Chevrier *et al.*, 2017, 2018). Cell-type assignment was performed using a Random Forest classifier trained on multiple manually gated samples. To investigate the utility of SCIM, we considered a subset comprising B-Cells and T-Cells only, for a total of *n*=135.334 cells. This dataset is further referred to as *target* dataset.

#### 3.2.2 scRNA Data Preparation

A second aliquot of the same patient sample was analyzed by droplet-based scRNA-sequencing using the 10x Genomics platform. A detailed description of the data analysis workflow is beyond the scope of this work and will be published elsewhere. In brief, standard QC-measures and preprocessing steps, such as removal of low quality cells, as well as filtering out mitochondrial, ribosomal and non-coding genes, were applied. Expression data was library-size normalized and corrected for the cell-cycle effect. Cell-type identification was performed using a set of cell-type-specific marker genes (Tirosh et al., 2016). Genes were then filtered to retain those that could code for proteins measured in CyTOF channels, the top 32 T-Cell/ B-Cell marker genes, and the remaining most variable genes for a final set of 256. The total number of B-Cells and T-Cells in this dataset amounts to *m*=4.683. The scRNA dataset is used as *source* dataset throughout the manuscript.

**Table 1:** Characteristics of the dataset derived from the melanoma sample.

| Dataset | # Markers | # Cells | T Cells | B Cells |
| --- | --- | --- | --- | --- |
| CytoF | 41 | 135.334 | 70% | 30% |
| scRNA | 1.024 | 4.683 | 73% | 27% |

### 3.3 Single-cell Profile of Human Bone Marrow

Oetjen *et al.* (2018) used several bulk and single-cell technologies to comprehensively characterize human bone marrow. The data was obtained from 20 healthy donors, whereas all data modalities were acquired for 8 samples. For our application we consider the single-cell transcriptome profile as well as CyTOF measurements of sample *O* from this dataset, that were carried out with the objective of describing in detail a T-Cell population. The data was preprocessed as described in Oetjen *et al.* (2018). The cell-type information for scRNA data was obtained directly by the courtesy of the authors, whereas CyTOF cells were manually gated using the strategy presented in **Supplementary Figure S7**. A subpopulation of CD8 naive T-Cells was filtered out due to a very small number of cells. The preprocessed data of the analysed sample included several T-Cell subtypes.

**Table 2:** Characteristics of the preprocessed dataset derived from the human bone marrow sample.

| Dataset | # Markers | # Cells | CD8 effector | CD4 naive | CD4 memory |
| --- | --- | --- | --- | --- | --- |
| CytoF | 34 | 98.799 | 57% | 26% | 17% |
| scRNA | 256 | 2.388 | 46% | 31% | 23% |

## 4 Experiments

### 4.1 Three Technology Simulated Data

We apply SCIM to integrate the three simulated datasets. The discriminator is trained to classify the source technology and is fully supervised using the branch label. The latent space is intialized by training a VAE on the source technology. The latent representations of the source technology are fixed, and the two target technologies are trained for 256 epochs. Bipartite matching is performed for each pair of datasets, using *k* = 64 and a null match penalty set to the 95^*th*^ percentile of edge costs.

### 4.2 Integration of scRNA and CyTOF Patient Data

We apply SCIM to integrate two sets of scRNA/CyTOF data, one set corresponds to a melanoma tumor from the Tumor Profiler project (Irmisch *et al.*, 2020) and the other one to a human bone marrow sample from Oetjen *et al.* (2018). The scRNA technology was chosen both times as the source technology, and the latent space is initialized by training a VAE. SCIM is trained for 64 epochs to integrate the CyTOF technologies. The discriminator is trained in both cases to classify the source technology and supervised with cell type labels. In the Bone Marrow cohort, the discriminator is fully supervised with the full cell type label. This was necessary as the number of available scRNA cells was quite small. For the Melanoma dataset we employed a semi-supervised strategy and use only 10% of the cell type labels. Bipartite matching is performed in both cases using *k* = 50 and a null match penalty set to the 95^*th*^ percentile of edge costs. In both scenarios, CyTOF cells greatly out number the scRNA cells, thus many-to-one scRNA matches are allowed.

## 5 Results

We evaluate the SCIM framework on a simulated dataset based on PROSSTT (Papadopoulos *et al.*, 2019) as well as two real world settings, where we match cells from CyTOF and scRNA measurements taken from a single sample analyzed within the Tumor Profiler project (Irmisch *et al.*, 2020) and from a human bone marrow sample Oetjen *et al.* (2018). We provide an implementation of the proposed approach in python using TensorFlow (Abadi *et al.*, 2015).

### 5.1 SCIM Aligns Substructure in Simulated data

Branches in PROSSTT define an over-arching structure that mimics cell-types, while the temporal component, i.e., pseudotime, provides a continuous interpolation from one branch to another as described by the tree (**Figure 3**). In latent space, the branch structure within the data produces large clusters, while the pseudotime structure provides orientation within each cluster as well as a globally smoothing of the manifold. SCIM is run to integrate the three simulated datasets producing a technology-invariant latent space (see **Figure 4**). SCIM embeddings capture the branching process and furthermore correctly orients the substructure of most branches (see **Figure 3**). We report 86% of matches retained the branch label and strong correlations among pseudotime (Pearson: 0.86, Spearman: 0.83) using a null node penalty of 95^*th*^ percentile that controls the false/true positive trade-off (see **Supplemental Figure S3**). Furthermore, most branch mismatches occur at the nodes of the tree, where the label is ambiguous due to the continuous nature of the temporal process (see **Supplemental Table S4**).

**Figure 4:**
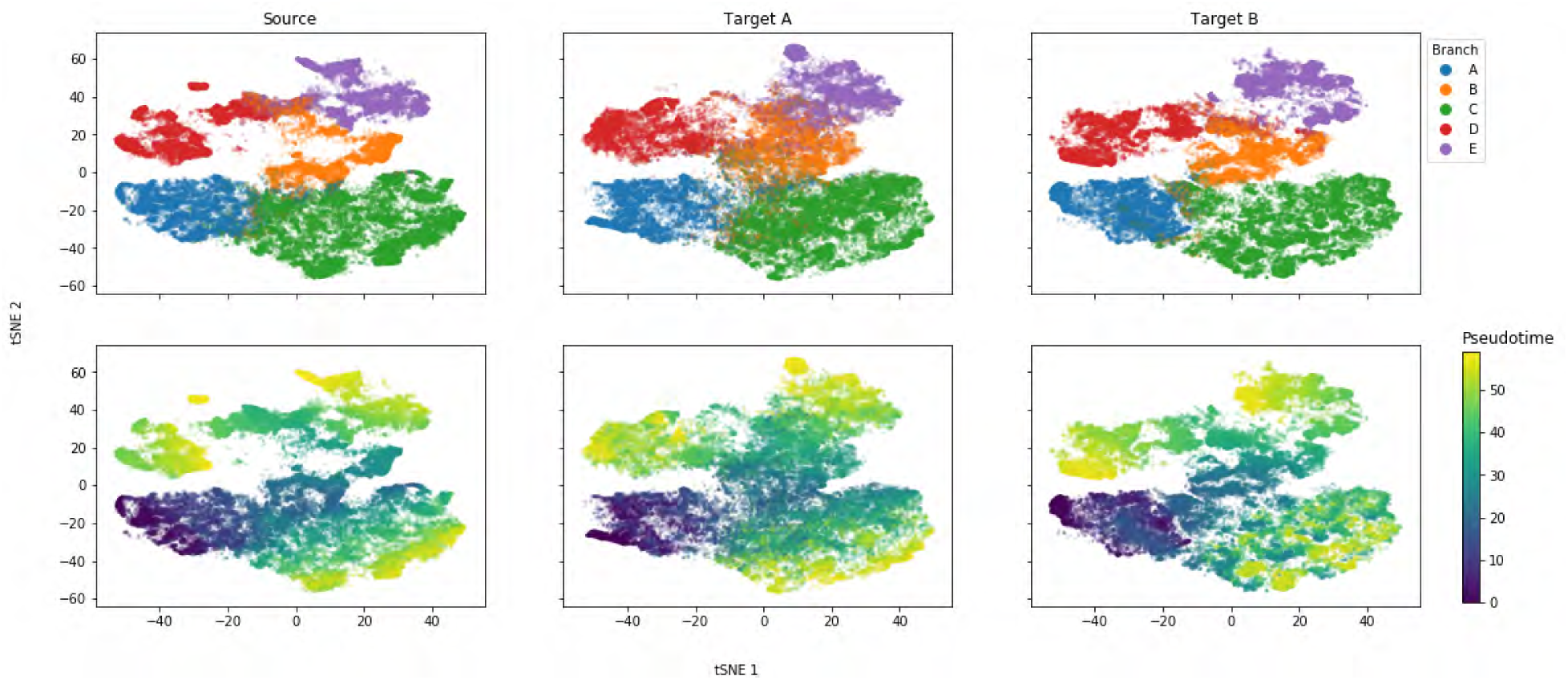
Integrated latent space of three synthetic datasets. Three single-cell ‘omics datasets are generated (Papadopoulos *et al.*, 2019) from a shared underlying temporal branching process (as defined in **Figure 3**). The same branching process was used in all three cases (as can be seen in the top row), but the parameters governing their feature distributions are drawn with different seeds. Hence, their latent structure is the same, yet they share no correspondences between features. SCIM is run, fully supervised using the branch label, and all datasets are embedded into a shared latent space. tSNE embeddings (Maaten and Hinton, 2008) are computed and visualized on the combined latent representations from all three datasets. Each column shows only the cells from a single technology. Cells are colored by their branch and pseudotime label in the first and second row, respectively.

The SCIM framework can be applied to a many-technology setting, and we demonstrate this by obtaining pairwise matches between all three datasets. SCIM successfully aligns the cells, based on evaluations on pseudotime (see **Supplemental Figure S1**) as well as branch label (see **Supplemental Table S5 and S6**), even when using codes from such an extended latent space.

These results demonstrate that SCIM is capable of accurately identifying the best matching cells across multiple technologies, based on the shared latent representations in the presence of an underlying branching process but in the absence of paired features.

#### 5.1.1 MATCHER Comparison: Capturing Complex Latent Structure

We compare SCIM to MATCHER (Welch *et al.*, 2017), which is, to the best of our knowledge, the only other published work that can integrate multi-modal ‘omics datasets in the absence of direct feature correspondences. MATCHER, however, assumes a one-dimensional latent structure that cannot capture hierarchical relationships, such as the ones exhibited in the simulated PROSSTT data, and frequently found and studied in single-cell datasets. Moreover, MATCHER is built around a GPLVM (Lawrence, 2004), which limits its scalability. To this end, we set a budget of 48 hours compute time and limit memory consumption to 40 Gb. Using the latent representations generated by MATCHER, we solve the bipartite matching problem setting the same hyperparameter configuration. MATCHER is unable to model the PROSSTT branching structure and is outperformed by SCIM with respect to matching (see **Supplemental Table S7, Supplemental Table S8**, and **Supplemental Figure S2**).

### 5.2 Universal Divergence Scales Model Selection in SCIM

To evaluate the performance of SCIM on real data and to gain a better understanding of the individual components of our framework, we apply SCIM on a melanoma tumor sample from the Tumor Profiler Consortium (Irmisch *et al.*, 2020). Model selection in the adversarial setting with real-world data is challenging since there is no metric that captures model performance, nor does one have access to any ground truth data to evaluate on. To help model selection, we employ a universal divergence estimator (Wang *et al.*, 2009) to evaluate the quality of the integrated latent space (see **Methods, Supplemental Figure S6**). This score measures how well two sets of points are mixed, and it is computed pairwise between source and target technologies. An optimization is defined as a success if the divergence and reconstruction errors are below the empirically set thresholds. This allows the evaluation of many model settings at scale despite operating in the adversarial setting. We find that performance depends on tuning *β* and the learning rates for the discriminator and encoder/decoder networks (see **Supplemental Table S3**).

### 5.3 Modified Bi-partite Matching is Scalable

Due to the large number of cells profiled with each individual technology per sample, we precede our bipartite matching with a kNN search (see **Section 2.2**). This reduces the problem complexity by *a priori* discarding redundant edges in the graph. Experiments on the real-world melanoma sample investigating the level of sparsity, governed by a hyperparameter *k*, show that using even a small number of neighbors provides good matching accuracy and performance saturates past *k* > 100 (see **Supplemental Figure S4, Supplemental Table S2**). This is in line with our expectations since a match to an extremely distant neighbor is hard to justify. In order to maintain a high degree of sparsity, without sacrificing matching accuracy, we use *k* = 50 in all further experiments.

### 5.4 SCIM Pairs Cells Across scRNA and CyTOF in a Melanoma Sample

Integrating data from scRNA and CyTOF technologies applied to a melanoma sample allows for a multi-view perspective on cell dynamics and, thus, will eventually lead to a more thorough understanding of the undergoing biological processes. Therefore we have evaluated the aforementioned melanoma sample with the SCIM framework. The bipartite matching on the latent codes has a 93% accuracy in recovering the cell-type label, calculated as the fraction of True Positives over all matches. A more fine-grained visual evaluation is performed by inspecting the matches on a tSNE embedding of the integrated latent space marked by grey lines (see **Figure 5**). The latent representation is explored thoroughly as 99.7%, and 99.3% of cells are matched to their analogs, from CyTOF and scRNA datasets, respectively. Given different cell-type proportions in the data, a certain number of mismatches is expected, which corresponds to the lines joining points across the two cell types. In comparison, a simple data-space matching approach would only utilize 5% of the scRNA cells (see **Supplemental Table S1 and Supplemental Figure S5**). To evaluate the latent space matching further, we used a more fine-grained information of marker expression correlation, to quantitatively assess the latent-space matching quality. We used the correlation coefficients between the expression of immuno-markers *CD20* and *CD3*. Both markers are characteristic to a subset of our data, as they are used to differentiate B-Cells and T-Cells, respectively. We found that matching using shared latent representations provides relatively high correlation coefficients (Pearson’s coefficient: 0.68 *CD20* and 0.55 for *CD3* marker, see **Figure 6**), given the expected low correlation between RNA expression and protein abundance. In conclusion, even in the presence of a subset of paired features across the technologies, using the shared latent representations proves beneficial for finding cell analogs.

**Figure 5:**
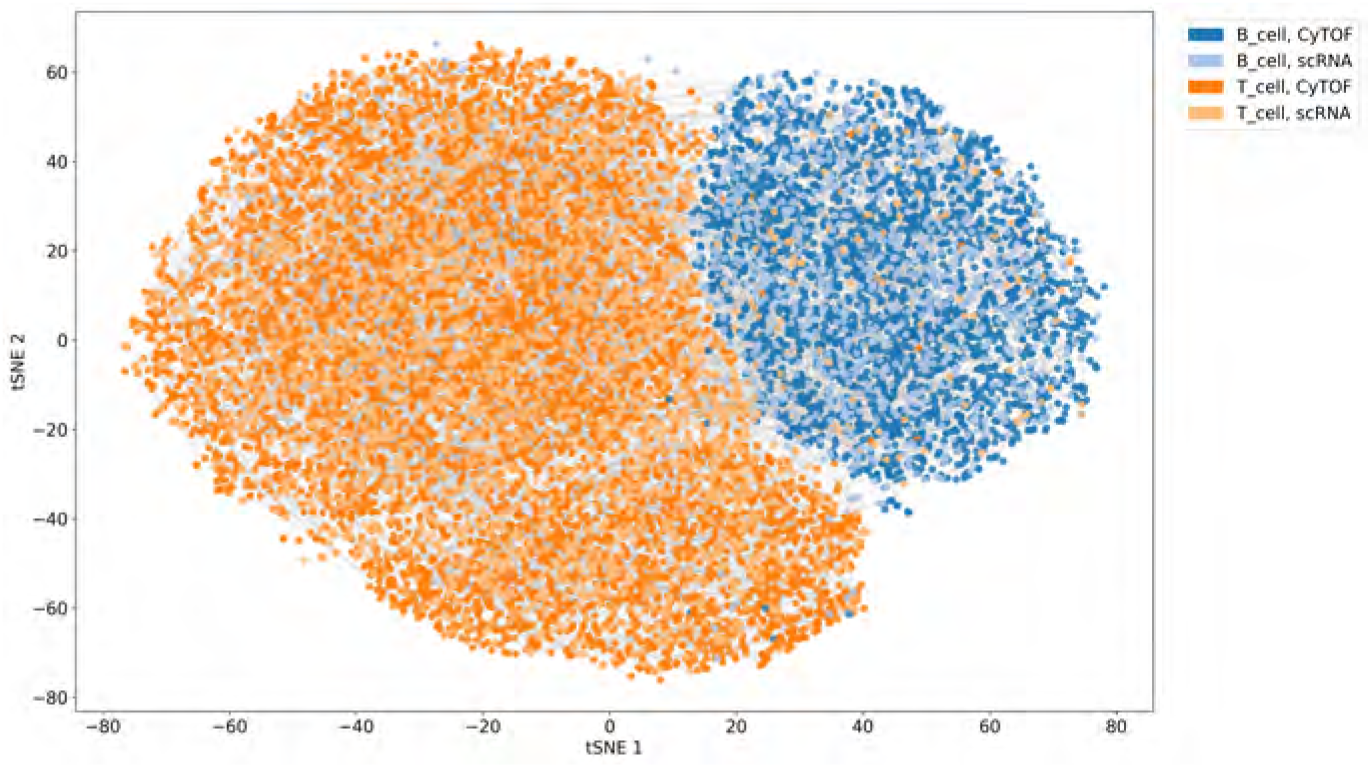
Integrated latent space and matches of scRNA and CyTOF cells from a melanoma sample from the Tumor Profiler Consortium. Discriminators are semi-supervised using 10% of the cell-type labels. Cells are colored by their cell-type label and shaded by their technology (dark shades: CyTOF, light shades: scRNA). Matches produced by SCIM are represented by grey lines connecting cells. tSNE embeddings (Maaten and Hinton, 2008) are computed on the whole dataset and then 10.000 matched pairs are sampled at random for visualization.

**Figure 6:**
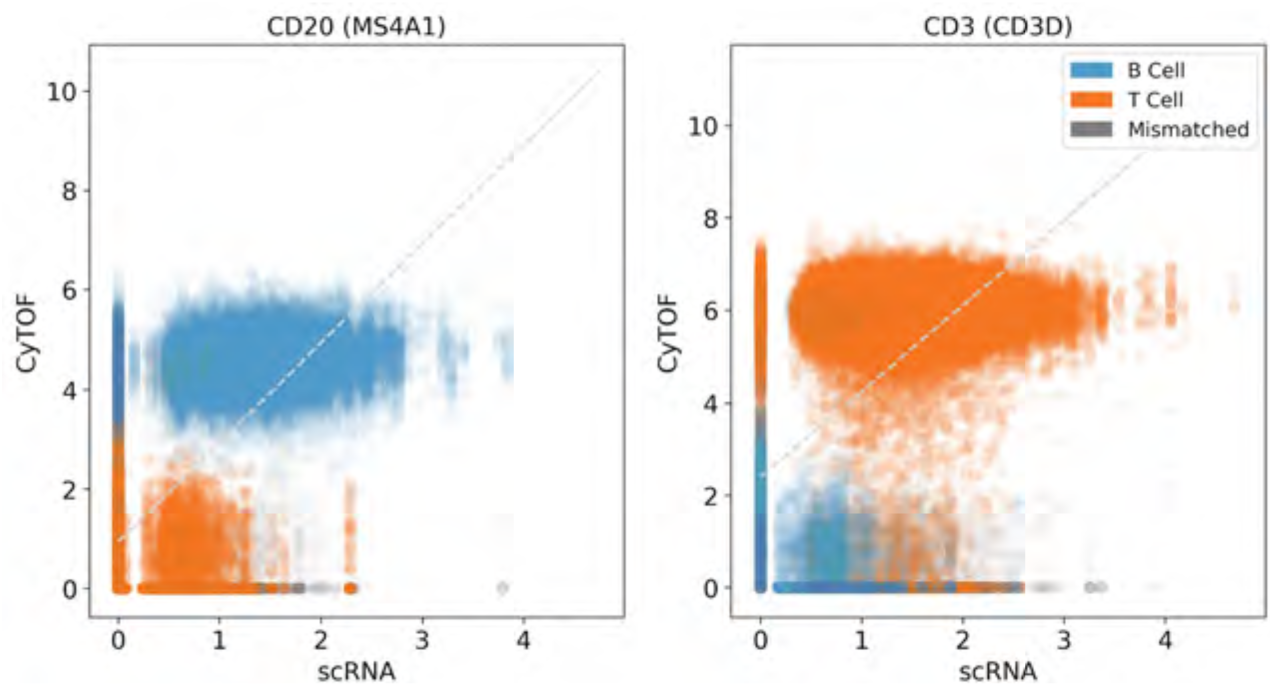
*CD20* and *CD3* marker abundances measured with scRNA (gene, x-axis) and CyTOF (protein, y-axis) in a melanoma sample from the Tumor Profiler Consortium. The values on the axes represent normalized expression. Colors (blue, orange) represent cell-types (B-Cell, T-Cell) while grey marks mismatches with respect to the cell-type label. The linear regression line is depicted by a dashed light grey line. Pearson’s correlation coefficient equals 0.68 and 0.55 for CD20 and CD3, respectively. Spearman’s correlation coefficient amounts to 0.6 and 0.45., for CD20 and CD3, respectively.

### 5.5 SCIM Recovers T-CellsSubpopulations Across Multi-Modal Human Bone Marrow Data

We use SCIM to integrate T-Cells derived from one sample in the Human Bone Marrow study, profiled with scRNA and CyTOF technologies. The tSNE embedding based on the latent space codes implies good integration across technologies while preserving the cell-subtype structure **Supplemental Figure S8**. We evaluate the quality of matches using fine-grained labels indicating one of the T-Cell subtypes identified in gating: CD8 effector, CD4 naive, CD4 memory (see **Section 3.3**). In a fully supervised approach, using the labels to orient the latent space, we achieve an accuracy of 88% with less than 1% of matches directed to the null node. Nevertheless, when utilizing only 10% of the labels in the semi-supervised approach, we note only a slight drop in performance, obtaining 84% correct matches with less than 1% of cells directed to the null node. Evaluating on higher-level labels of CD8 vs. CD4 T-Cells improves the accuracy to 96% and 91% for the fully supervised and semi-supervised approach, respectively. As expected, distinguishing cellular subtypes (e.g., CD4 naive versus CD4 memory) is more challenging due to high overall similarity between the cell populations, but overall SCIM is capable of accurately recovering even such subtle differences between cell type and states.

## 6 Discussion

We have developed SCIM, a new technology-invariant approach that pairs single-cell measurements across multimodal datasets, without requiring feature correspondences. This development enables real multi-modal single-cell analysis, and opens up new opportunities to gain a multi-view understanding of the dynamics of individual cells in various disease or developmental states. The underlying auto-encoder framework, combined with a customized bipartite matching approach, ensures scalability even with large numbers of cells.

We demonstrate that our model performs well on simulated data as well as on real-world melanoma and bone marrow samples profiled with scRNA and CyTOF. The SCIM framework is presented here on two and three data modalities and easily extends to additional technologies, providing a new and effective solution to the multi-level data integration problem. Integration of Image Mass Cytometry (Giesen *et al.*, 2014) or single-cell ATAC-seq data (Buenrostro *et al.*, 2015) for example, could enable the spatial analysis of regulatory and global expression changes not just in cancer but also in other diseases such as multiple sclerosis, where detailed spatiotemporal information has already shown to provide relevant insights (Ramaglia *et al.*, 2019). Notably, SCIM allows for the integration of cell populations undergoing branching processes, enabling the study of temporal phenomena, such as developmental and cell fate determination. The scalability of our framework ensures applicability beyond single samples, facilitating the study of large cohorts or the integration of SCIM into analytical workflows.

As the low-dimensional representation of the data produced by SCIM is used solely to perform cell matching, it can be combined with any other analytical methods. The truly observed signals per cell pair measured with different technologies can be used for any downstream analysis. By adopting a divergence measure (Wang *et al.*, 2009), we addressed common constraints in adversarial training, such as training instability and convergence problems. To ensure scalability, we used a modified bipartite matching solution to efficiently match corresponding cells across technologies. Our extensions guarantee wider applicability of SCIM, since shifts in cell-type composition across disjoint aliquots, even coming from the same sample, can be expected. Furthermore, the introduction of the *null* node ensures a higher quality of matches by avoiding forced mismatches and thus, improving confidence in the cell-to-cell assignments. With increasing data dimensionality, the number of nearest neighbours (*k*) should also rise, since more ties are likely to occur. Nevertheless, the difference in the number of True Positives across various values of *k* for the same dataset remains within 5% in our experiments. Hence, we can state that performance is robust against the choice of this hyperparameter. Furthermore, SCIM is itself inherently modular, and other matching strategies that may be more suitable to other data types or experimental designs can be easily deployed on the integrated latent codes.

SCIM helps bridge the gap between data generation and integrative interpretation of diverse multi-modal data in the rapidly expanding field of single-cell biology, providing users with an easily scalable algorithm designed to maximize the information it provides and not limited to fit a particular analytical approach.

## Acknowledgments

We would like to thank Christopher S. Hourigan and Gege Gui for providing us with the cell-type information for the scRNA dataset from Oetjen *et al.* (2018). Furthermore, we would like to thank Felix Faltings for pointing us to the PROSSTT simulation framework and for his help with the set-up.

## Funding

This project is jointly funded by a public-private partnership involving Roche, ETH Zürich, University of Zürich, University Hospital Zürich, and University Hospital Basel. Francesco Locatello is supported by the Max Planck ETH Center for Learning Systems, by an ETH core grant (to Gunnar Rätsch), and by a Google Ph.D. Fellowship.

## 7 Supplement

**Figure S1:**
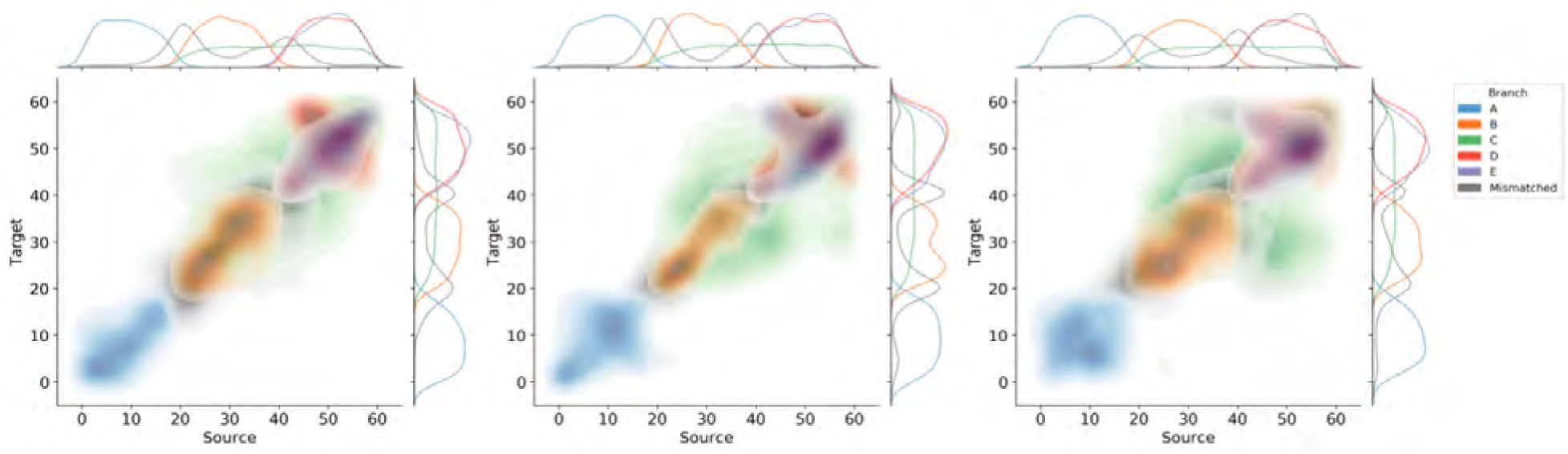
Evaluation of cross-technology cell matches made by SCIM on the simulated data with three technologies. The pairwise matching is attained for Source-Target A, Source-Target B, and Target A-Target B, respectively. Here, we show a density plot for matched pseudotime values between the source technology and the target technology, colored by the branch label. Mismatched cells are colored in grey. The tree defining the temporal branching process can be found in Figure 4, left. Marginal distributions of cell pseudotime for each branch is shown on the top and right.

**Figure S2:**
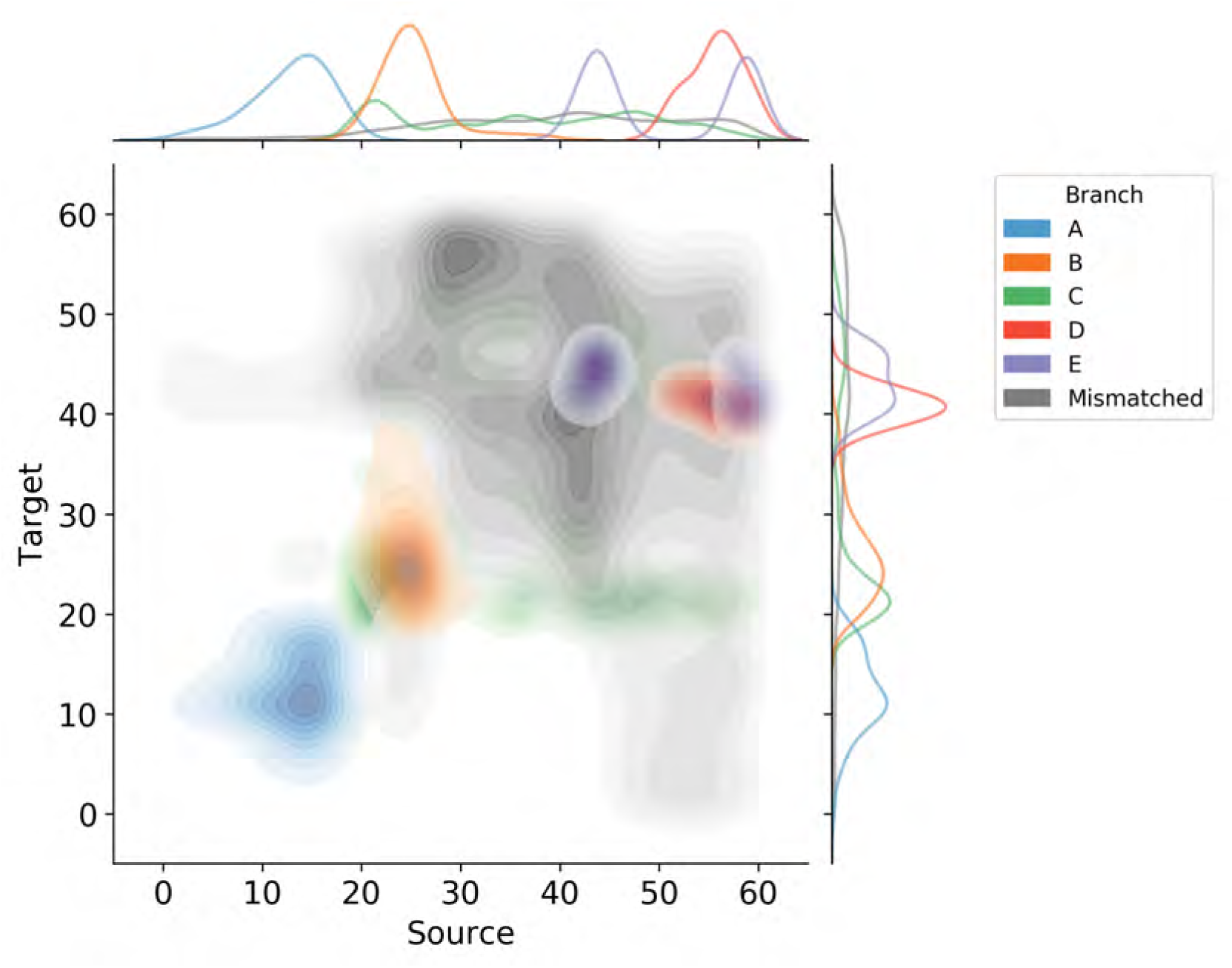
Evaluation of cross-technology cell matches using latent representation obtained by MATCHER on the simulated data. Cells are matched across datasets pairwise using the bipartite matching scheme. Here we show a density plot for matched pseudotime values between the source technology and the target technology, colored by the branch label. Mismatched cells are colored in grey. Marginal distributions of cell pseudotime for each branch are shown on the top and right.

**Figure S3:**
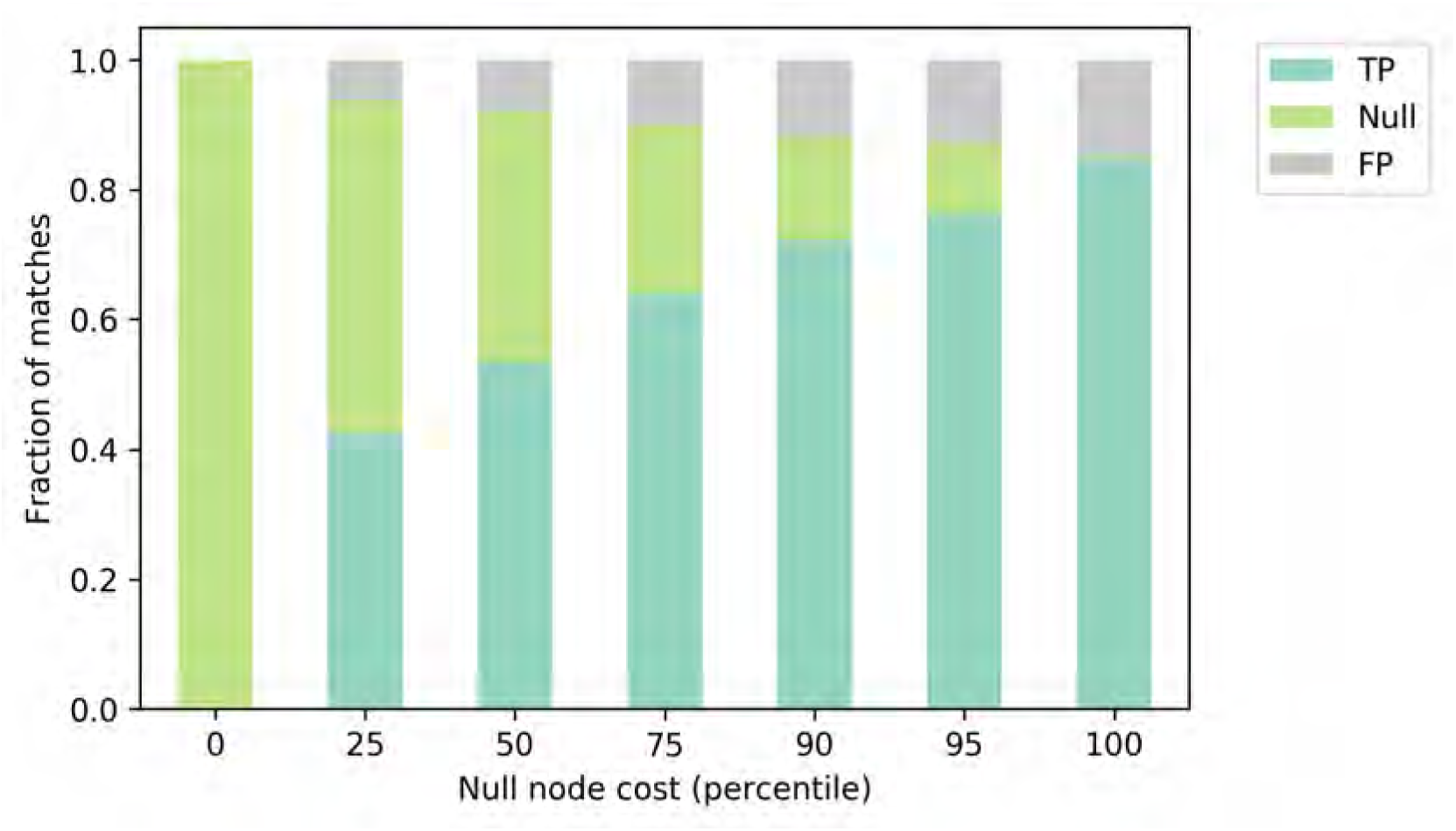
True Positive, False Positive, and Null matches fraction of all the matched pairs in the PROSSTT Source-Target experiment as a function of the cost for matching to the null node. The cost is specified as the percentile of the costs on all the other edges in the graph, i.e., Euclidean distance between latent codes of cells from source and target technologies. With decreasing null node cost, more False Positives are removed, on the trade-off of removing more True Positives.

**Figure S4:**
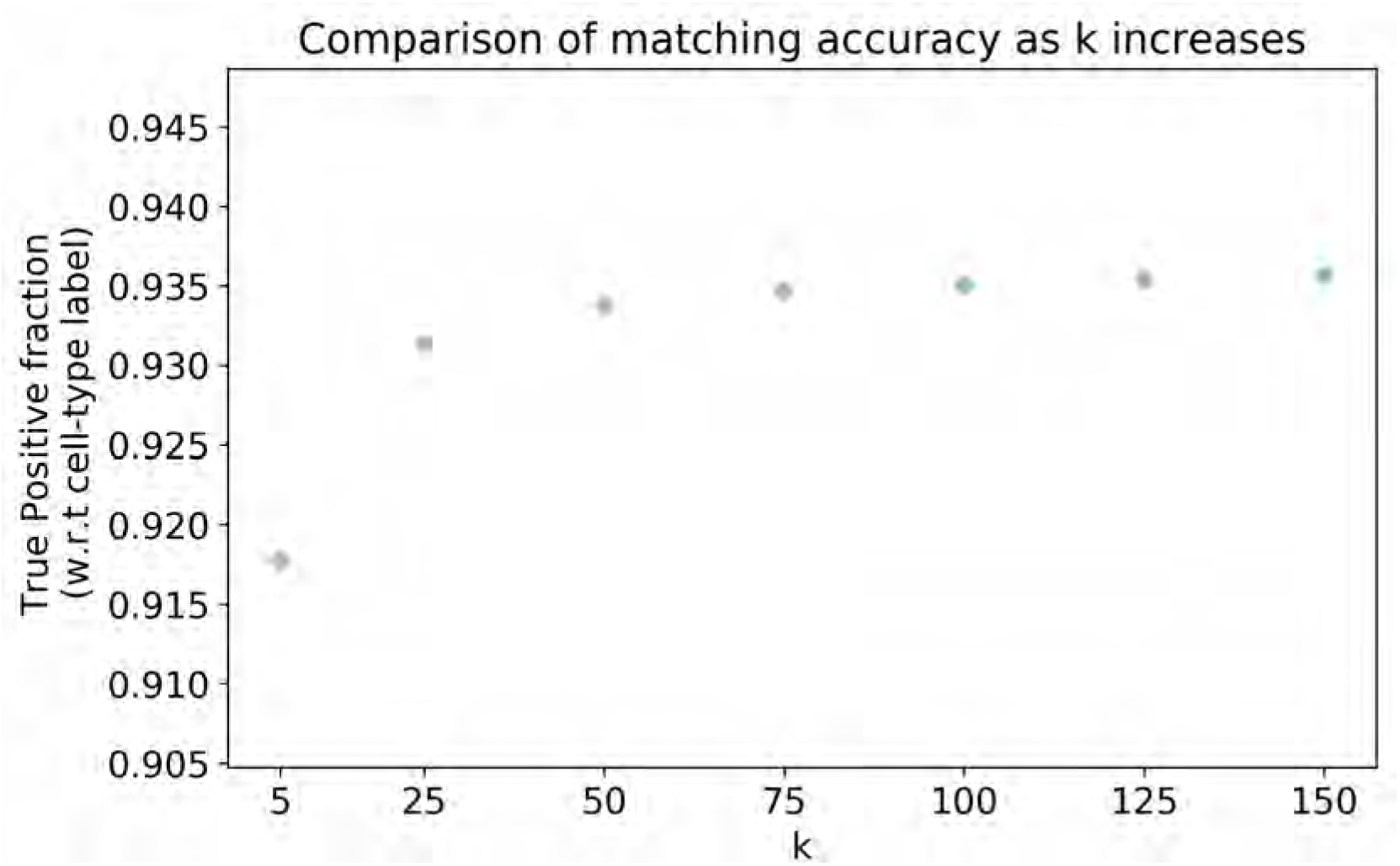
Comparison of matching accuracy, with respect to cell-type label as less sparsity of connections, is imposed on the Tumor Profiler data. The number of considered Nearest Neighbors *k* is indicated on the x-axis, and the fraction of True Positive matches, with respect to cell-type label, is depicted on the y-axis. The accuracy level saturates with *k* = 100.

**Figure S5:**
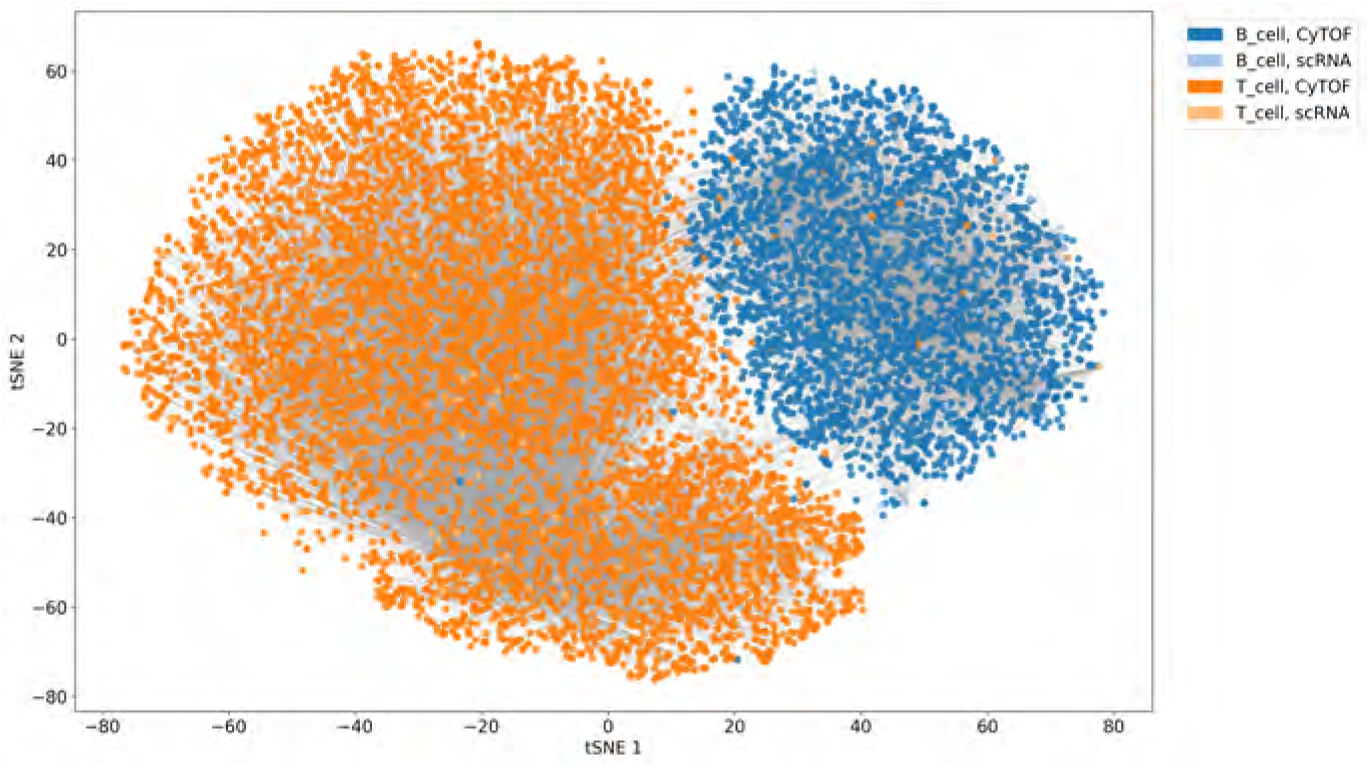
Matches of scRNA and CyTOF cells from the melanoma patient computed on raw expression values. The latent space is constructed as in **Figure 5**, but matches are made using distances computed between the expression of 37 genes and proteins that could share a correspondence between datasets.

**Figure S6:**
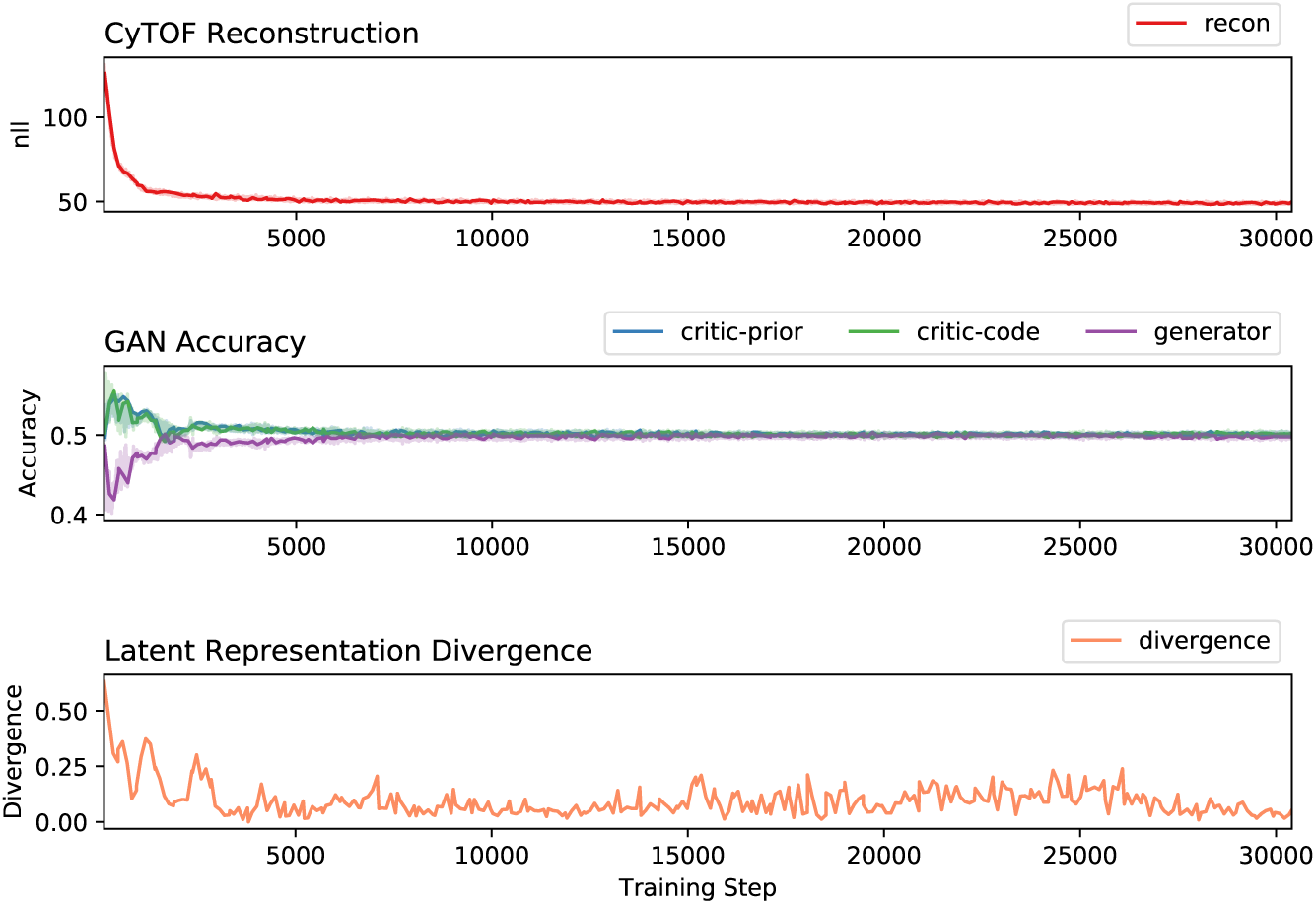
Training progress of SCIM on a melanoma sample. The latent space is initialized by training a VAE on scRNA data. SCIM integrates CyTOF representations into the latent space defined by the scRNA codes. The top panel shows the negative log-likelihood of the CyTOF reconstruction. The middle panel shows the performance of the discriminator to correctly classify scRNA codes (critic-prior), CyTOF codes (critic-code), and the ability of the encoder to fool the discriminator, i.e., the misclassification accuracy (generator). The bottom panel shows the divergence of the latent representations. The model is able to converge quickly. However the divergence score can fluctuate despite still fooling the discriminator. Training took 7815 seconds (just over 2 hours) and had a peak memory consumption of 1068 MB.

**Figure S7:**
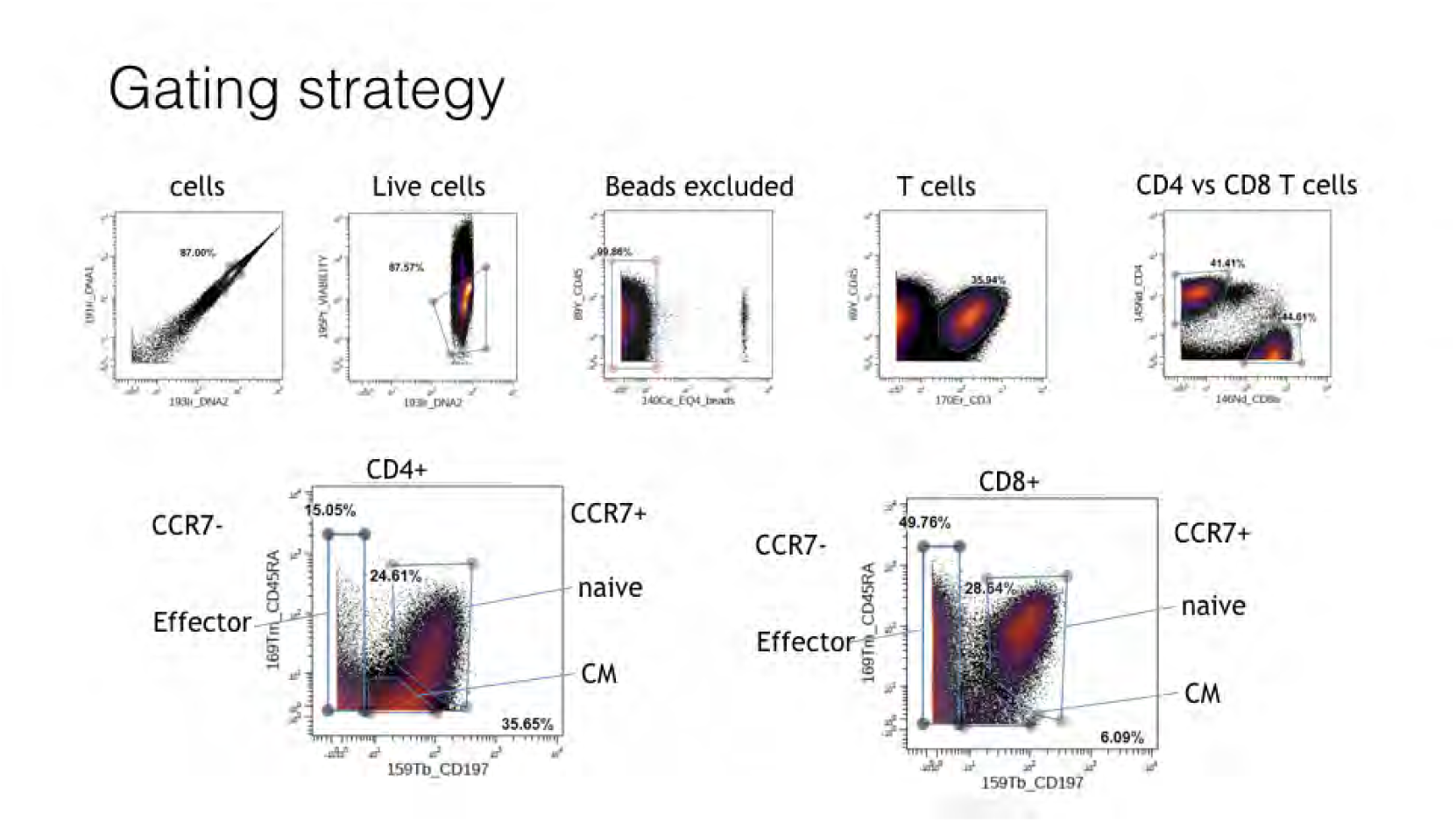
Gating strategy for cells profiled with CyTOF from the bone marrow patient.

**Figure S8:**
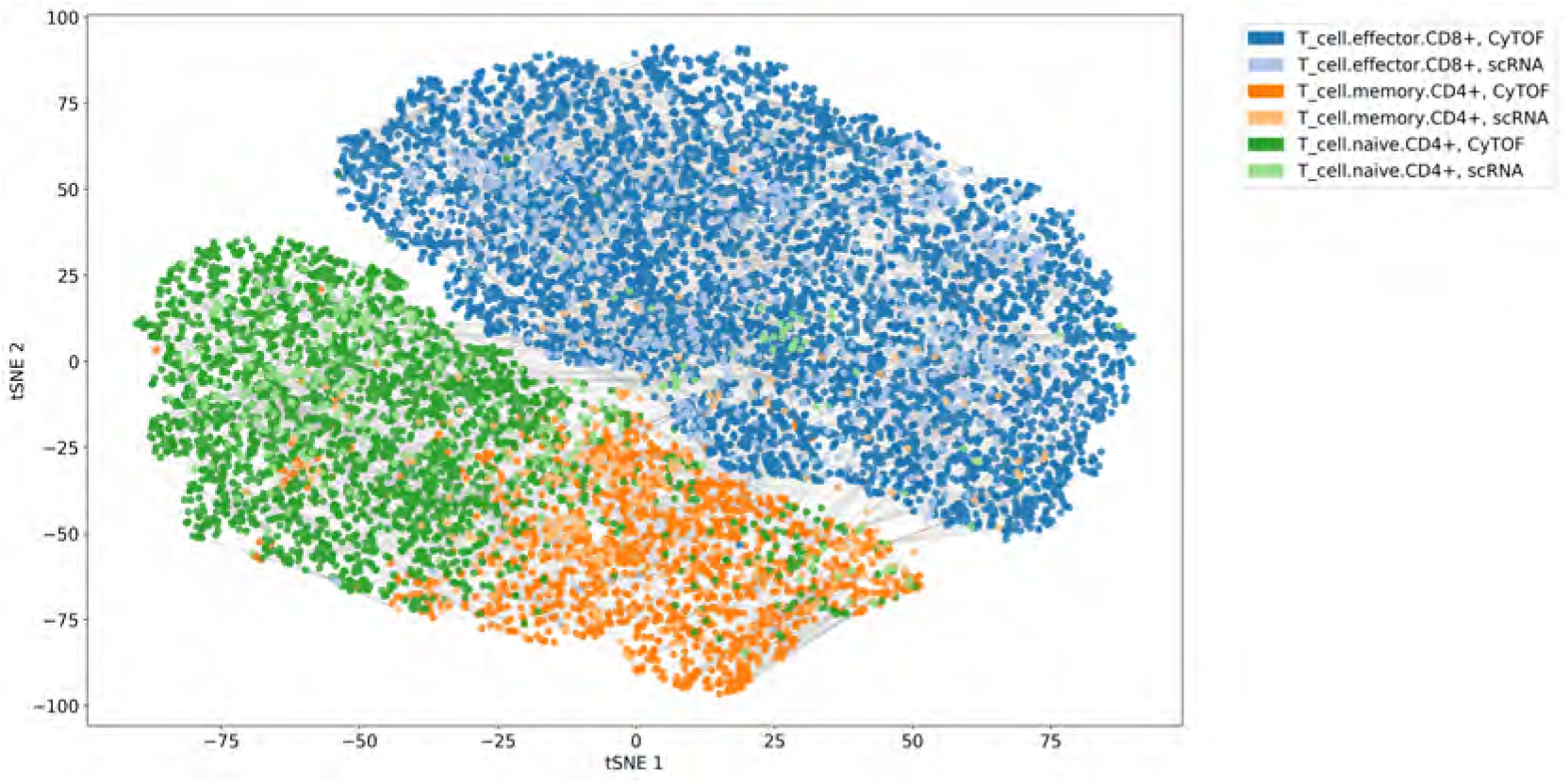
A tSNE embedding (perplexity=30) of the integrated latent space with cell matches indicated by the grey lines. The shared representation was obtained in a semi-supervised fashion, utilizing 10% of the cell-type labels to orient the latent space. 10.000 matched pairs were sampled at random for the plot. Colors (blue, green, orange) represent T-Cellsubtypes (CD8 effector, CD4 naive, CD4 memory), and color shades correspond to the profiling technology (light: scRNA, dark: CyTOF).

**Table S1:** The fraction of the cells from the source (scRNA) and target (CyTOF) datasets in TuPro that are matched using the Minimum-Cost Maximum-Flow algorithm. The matching is performed using the shared latent codes or the corresponding features in the data space. The data-space matching results in all the matches collapsing onto very few cells (5% of the source dataset). Using latent codes allows for exploration of the whole space and providing optimal matches for almost all the cells (less than 1% left unmatched).

| space | #cells matched<br>(source) | fraction<br>(source) | #cells matched<br>(target) | fraction<br>(target) |
| --- | --- | --- | --- | --- |
| latent | 4652 | 0.993 | 134930 | 0.997 |
| data | 236 | 0.05 | 130899 | 0.967 |

**Table S2:** Memory usage and computation time of the optimal matching as *k* hyperparameter in kNN search increases. The values were obtained on the whole TuPro dataset using the MCMF algorithm on an extended graph, as described in section 2.2.3. The cost of matching to the null node was set to 95th percentile, and a union of connections obtained from kNN graphs with *k* indicated in the first column was used for matching. The memory and time reports were averaged across three independent runs.

| k | CPU time [s] | Max Memory [Mb] |
| --- | --- | --- |
| 5 | 547.00 | 1674.16 |
| 15 | 1599.00 | 3131.53 |
| 25 | 2940.67 | 4691.31 |
| 50 | 7015.33 | 8745.39 |
| 75 | 12306.33 | 12357.72 |
| 100 | 17875.67 | 17019.24 |
| 125 | 17200.00 | 20618.30 |
| 150 | 22286.00 | 24226.19 |

**Table S3:** Ablation study for training SCIM on the Melanoma patient at several levels of semi-supervision. Fraction censored is the fraction labels removed during training. The top 4 configurations for each level of semi-supervision is shown. *β* is the regularization strength of the adversarial loss. Learning rates are the initial settings of the ADAM optimizer. If the latent space divergence is below 0.3 and the negative log-likelihood of the input under the reconstruction is below 47, the training is determined to be a success. These values were chosen empirically. We see that *β* and optimizer learning rates were heavily influential on model success.

| Fraction Censored | $\beta$ | Encoder/Decoders | Critic | Success Rate | Total Runs |
| --- | --- | --- | --- | --- | --- |
| 0.00 | 16 | 0.0010 | 0.0010 | 14% | 7 |
|  | 16 | 0.0010 | 0.0005 | 14% | 7 |
|  | 16 | 0.0005 | 0.0005 | 10% | 10 |
|  | 16 | 0.0005 | 0.0010 | 10% | 10 |
| 0.25 | 16 | 0.0010 | 0.0010 | 14% | 14 |
|  | 16 | 0.0005 | 0.0010 | 12% | 17 |
|  | 64 | 0.0001 | 0.0005 | 0% | 7 |
|  | 64 | 0.0001 | 0.0010 | 0% | 7 |
| 0.75 | 16 | 0.0005 | 0.0010 | 17% | 6 |
|  | 32 | 0.0005 | 0.0005 | 12% | 8 |
|  | 16 | 0.0010 | 0.0005 | 8% | 13 |
|  | 64 | 0.0001 | 0.0005 | 0% | 7 |
| 0.90 | 16 | 0.0005 | 0.0010 | 40% | 10 |
|  | 16 | 0.0010 | 0.0005 | 10% | 10 |
|  | 16 | 0.0005 | 0.0005 | 8% | 12 |
|  | 32 | 0.0010 | 0.0001 | 0% | 10 |
| 0.95 | 32 | 0.0010 | 0.0001 | 0% | 4 |
|  | 64 | 0.0001 | 0.0005 | 0% | 4 |
|  | 64 | 0.0001 | 0.0010 | 0% | 6 |
|  | 32 | 0.0010 | 0.0010 | 0% | 5 |
| 0.99 | 16 | 0.0005 | 0.0010 | 17% | 6 |
|  | 32 | 0.0010 | 0.0005 | 0% | 6 |
|  | 64 | 0.0001 | 0.0005 | 0% | 4 |
|  | 32 | 0.0010 | 0.0001 | 0% | 6 |

**Table S4:** Confusion table showing branch labels of the optimal matches of cells in the simulated PROSSTT data, between Source and Target A. Entries on the diagonal correspond to correct matches whereas off-diagonal elements to mismatches. The overall accuracy, with respect to the branch label, equals 86%.

| Target Source | A | B | C | D | E |
| --- | --- | --- | --- | --- | --- |
| A | 8528 | 583 | 679 | 99 | 23 |
| B | 724 | 6285 | 758 | 466 | 928 |
| C | 770 | 777 | 18917 | 48 | 99 |
| D | 42 | 371 | 22 | 7804 | 266 |
| E | 28 | 1086 | 41 | 305 | 7363 |

**Table S5:**
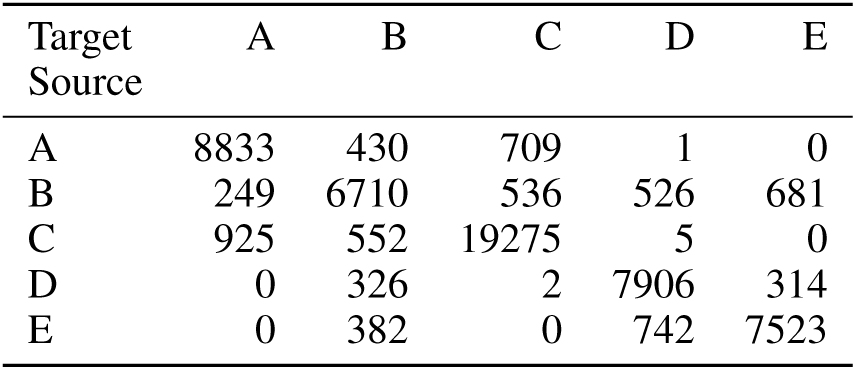
Confusion table showing branch labels of the optimal matches of cells in the simulated PROSSTT data, between Source and Target B. Entries on the diagonal correspond to correct matches whereas off-diagonal elements to mismatches. The overall accuracy, with respect to the branch label, equals 86%.

**Table S6:**
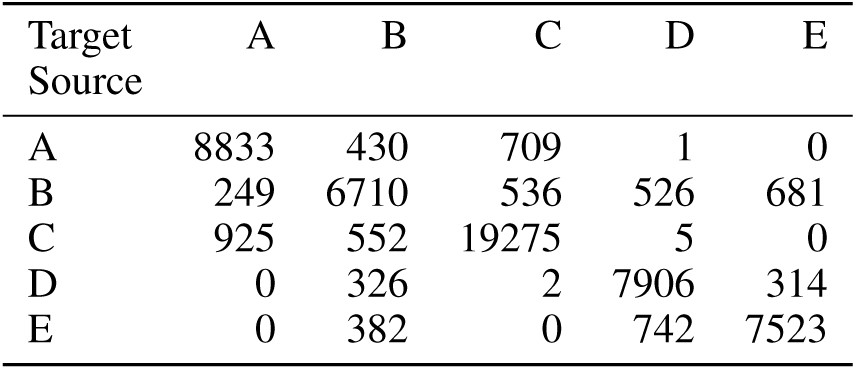
Confusion table showing branch labels of the optimal matches of cells in the simulated PROSSTT data, between Target A and Target B. Entries on the diagonal correspond to correct matches whereas off-diagonal elements to mismatches. The overall accuracy, with respect to the branch label, equals 84%.

**Table S7:**
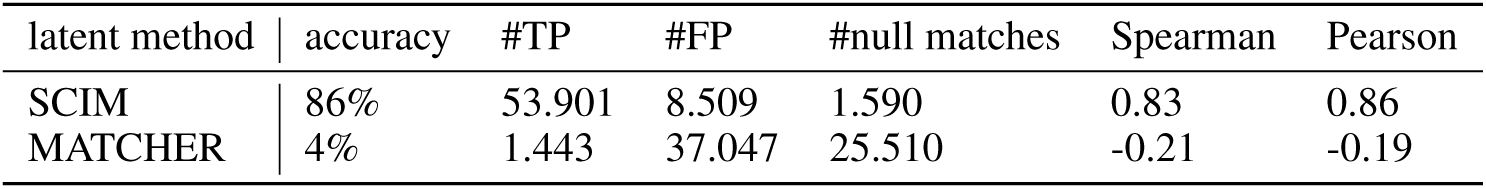
The matching results on the PROSSTT dataset, where the SCIM matching algorithm was applied to SCIM shared latent codes and the MATCHER latent representation. The table depicts accuracy with respect to branch label, the number of True and False positives, as well as null node matches and the correlation coefficients for the pseudotime between matched source and target cells.

**Table S8:** Confusion table showing branch labels of the matched cells in the simulated PROSSTT data. SCIM matching algorithm was applied to a shared latent representation obtained with MATCHER. Entries on the diagonal correspond to correct matches whereas off-diagonal elements to mismatches. The overall accuracy, with respect to the branch label, equals 4%.

| Target Source | A | B | C | D | E |
| --- | --- | --- | --- | --- | --- |
| A | 540 | 523 | 116 | 1133 | 453 |
| B | 713 | 788 | 5940 | 581 | 208 |
| C | 155 | 4998 | 100 | 4026 | 6526 |
| D | 173 | 274 | 5278 | 4 | 98 |
| E | 3162 | 203 | 2484 | 3 | 11 |

## TUPRO Consortium

Rudolf Aebersold (2), Faisal S Al-Quaddoomi (8,15), Jonas Albinus (7), Ilaria Alborelli (23), Per-Olof Attinger (10), Marina Bacac (14), Daniel Baumhoer (23), Beatrice Beck-Schimmer (29), Niko Beerenwinkel (4), Christian Beisel (4), Lara Bernasconi (26), Anne Bertolini (8,15), Bernd Bodenmiller (33), Ximena Bonilla (3,6,15,25), Ruben Casanova (33), Stéphane Chevrier (33), Natalia Chicherova (8,15), Maya D’Costa (9), Esther Danenberg (35), Natalie Davidson (3,6,15,25), Reinhard Dummer (27), Stefanie Engler (33), Martin Erkens (12), Katja Eschbach (4), Cinzia Esposito (35), André Fedier (16), Pedro Ferreira (4), Joanna Ficek (3,6,15,25), Anja L Frei (28), Bruno Frey (11), Sandra Goetze (7), Linda Grob (8,15), Detlef Günther (5), Pirmin Haeuptle (1), Viola Heinzelmann-Schwarz (16,22), Sylvia Herter (14), Rene Holtackers (35), Tamara Huesser (14), Anja Irmisch (27), Francis Jacob (16), Andrea Jacobs (33), Tim M Jaeger (10), Katharina Jahn (4), Alva R James (3,6,15,25), Philip M Jermann (23), André Kahles (3,6,15,25), Abdullah Kahraman (15,28), Viktor H Koelzer (28), Werner Kuebler (24), Jack Kuipers (4), Christian P Kunze (21), Christian Kurzeder (19), Jelena Kühn-Georgijevic (12), Kjong-Van Lehmann (3,6,15,25), Mitchell Levesque (27), Sebastian Lugert (9), Gerd Maass (11), Sergio Maffioletti (34), Julien Mena (2), Ulrike Menzel (4), Nicola Miglino (1), Emanuela S Milani (7), Holger Moch (28), Simone Muenst (23), Riccardo Murri (36), Charlotte KY Ng (23,32), Stefan Nicolet (23), Patrick GA Pedrioli (2), Lucas Pelkmans (35), Salvatore Piscuoglio (16,23), Michael Prummer (8,15), Mathilde Ritter (16), Christian Rommel (12), María L Rosano-González (8,15), Gunnar Rätsch (3,6,15,25), Jacobo Sarabia del Castillo (35), Ramona Schlenker (13), Petra C Schwalie (12), Severin Schwan (10), Tobias Schär (4), Gabriela Senti (26), Franziska Singer (8,15), Berend Snijder (2), Bettina Sobottka (28), Vipin T Sreedharan (8,15), Stefan Stark (3,6,15,25), Daniel J Stekhoven (8,15), Tinu M Thomas (3,6,15,25), Markus Tolnay (23), Nora C Toussaint (8,15), Mustafa A Tuncel (4), Audrey Van Drogen (7), Marcus Vetter (18), Tatjana Vlajnic (23), Sandra Weber (26), Walter P Weber (17), Rebekka Wegmann (2), Michael Weller (31), Fabian Wendt (7), Norbert Wey (28), Andreas Wicki (1,16,20), Bernd Wollscheid (7), Shuqing Yu (8,15), Johanna Ziegler (27), Marc Zimmermann (3,6,15,25), Martin Zoche (28), Gregor Zuend (30) (1) Cantonal Hospital Baselland, Medical University Clinic, Rheinstrasse 26, 4410 Liestal, Switzerland, (2) ETH Zurich, Department of Biology, Otto-Stern-Weg 3, 8093 Zurich, Switzerland, (3) ETH Zurich, Department of Biology, Wolfgang-Pauli-Strasse 27, 8093 Zurich, Switzerland, (4) ETH Zurich, Department of Biosystems Science and Engineering, Mattenstrasse 26, 4058 Basel, Switzerland, (5) ETH Zurich, Department of Chemistry and Applied Biosciences, Vladimir-Prelog-Weg 1-5/10, 8093 Zurich, Switzerland, (6) ETH Zurich, Department of Computer Science, Institute of Machine Learning, Universitätstrasse 6, 8092 Zurich, Switzerland, (7) ETH Zurich, Department of Health Sciences and Technology, Otto-Stern-Weg 3, 8093 Zurich, Switzerland, (8) ETH Zurich, NEXUS Personalized Health Technologies, John-von-Neumann-Weg 9, 8093 Zurich, Switzerland, (9) F. Hoffmann-La Roche Ltd, Grenzacherstrasse 124, 4070 Basel, Switzerland, (10) F. Hoffmann-La Roche Ltd, Grenzacherstrasse 124, 4070 Basel, Switzerland,, (11) Roche Diagnostics GmbH, Nonnenwald 2, 82377 Penzberg, Germany, (12) Roche Pharmaceutical Research and Early Development, Roche Innovation Center Basel, Grenzacherstrasse 124, 4070 Basel, Switzerland, (13) Roche Pharmaceutical Research and Early Development, Roche Innovation Center Munich, Roche Diagnostics GmbH, Nonnenwald 2, 82377 Penzberg, Germany, (14) Roche Pharmaceutical Research and Early Development, Roche Innovation Center Zurich, Wagistrasse 10, 8952 Schlieren, Switzerland, (15) Swiss Institute of Bioinformatics, Zurich, Switzerland, (16) University Hospital Basel and University of Basel, Department of Biomedicine, Hebelstrasse 20, 4031 Basel, Switzerland, (17) University Hospital Basel and University of Basel, Department of Surgery, Brustzentrum, Spitalstrasse 21, 4031 Basel, Switzerland, (18) University Hospital Basel, Brustzentrum & Tumorzentrum, Petersgraben 4, 4031 Basel, Switzerland, (19) University Hospital Basel, Brustzentrum, Spitalstrasse 21, 4031 Basel, Switzerland, (20) University Hospital Basel, Centre for Neuroendocrine & Endocrine Tumours, Spitalstrasse 21/Petersgraben 4, 4031 Basel, Switzerland, (21) University Hospital Basel, Department of Information- and Communication Technology, Spitalstrasse 26, 4031 Basel, Switzerland, (22) University Hospital Basel, Gynecological Cancer Center, Spitalstrasse 21, 4031 Basel, Switzerland, (23) University Hospital Basel, Institute of Medical Genetics and Pathology, Schönbeinstrasse 40, 4031 Basel, Switzerland, (24) University Hospital Basel, Spitalstrasse 21/Petersgraben 4, 4031 Basel, Switzerland, (25) University Hospital Zurich, Biomedical Informatics, Schmelzbergstrasse 26, 8006 Zurich, Switzerland, (26) University Hospital Zurich, Clinical Trials Center, Rämistrasse 100, 8091 Zurich, Switzerland, (27) University Hospital Zurich, Department of Dermatology, Gloriastrasse 31, 8091 Zurich, Switzerland, (28) University Hospital Zurich, Department of Pathology and Molecular Pathology, Schmelzbergstrasse 12, 8091 Zurich, Switzerland, (29) University Hospital Zurich, Institute for Anesthesiology, Rämistrasse 100, 8091 Zurich, Switzerland, (30) University Hospital Zurich, Rämistrasse 100, 8091 Zurich, Switzerland, (31) University Hospital and University of Zurich, Department of Neurology, Frauenklinikstrasse 26, 8091 Zurich, Switzerland, (32) University of Bern, Department of BioMedical Research, Murtenstrasse 35, 3008 Bern, Switzerland, (33) University of Zurich, Department of Quantitative Biomedicine, Winterthurerstrasse 190, 8057 Zurich, Switzerland, (34) University of Zurich, Grid Computing Competence Center, Rämistrasse 71, 8006 Zurich, Switzerland, (35) University of Zurich, Institute of Molecular Life Sciences, Winterthurerstrasse 190, 8057 Zurich, Switzerland, (36) University of Zurich, Services and Support for Science IT, Winterthurerstrasse 190, 8057 Zurich, Switzerland

## Notes

### Competing Interest Statement

The authors have declared no competing interest.

### Summary of Updates

update author metadata

